# Beyond Reach I: Neuroimaging Reveals Different Cortico-cerebellar Networks for Grasp vs. Object Placement

**DOI:** 10.64898/2026.04.21.719705

**Authors:** Gaelle N. Luabeya, Erez Freud, J. Douglas Crawford

## Abstract

Object manipulation usually involves both acquisition and placement, but neuroimaging studies tend to focus on its initial reach-to-grasp component. To understand how grasp and placement are represented in the brain, we designed an event-related fMRI study in which 20 participants alternated between grasping / placing a rectangular object from / toward templates presented at variable locations and orientations on an inclined plane. Despite the similarity in target information (i.e., location and orientation), we expected to see sensory and intentional task differences, as the grasp relied on matching the hand to the object, whereas placement relied on matching the object to the template. For our analysis, we examined BOLD activation across two epochs (planning phase before movement onset and execution phase during the movement), and applied graph-theoretical analysis (GTA) of whole-brain functional connectivity based on the joint time series from these two epochs. In both tasks, we found that common activation was more focused on prefrontal regions during planning and on sensorimotor regions during execution, with a specific increase in cerebellar activation during placement. Furthermore, the greater placement activation was also found in action-related regions of interest during the planning phase, and they could accurately decode between our two tasks. GTA categorized the two networks into three identical modules (prefrontal, cerebellar-occipito-parietal, and sensorimotor) and found task-based differences in modularity scores in the prefrontal and cerebellar-occipito-parietal modules. Overall, these data show that although reach-to-grasp and reach-to-place movements share extensive neural circuitry, they also exhibit task-specific differences, likely related to differences in intentionality during planning and in sensory feedback during movement execution.

## INTRODUCTION

Goal-directed actions, such as prehension, often begin with an initial reach-to-grasp movement, but this is usually just the prelude toward the ultimate intention: to manipulate and/or place the object somewhere. For example, when loading a dishwasher, although one first grasps a dirty plate, the real objective of the full movement is to place and orient the dish correctly in the lower rack and run a successful cleaning cycle. Such actions underscore the importance of considering not only movement initiation but also its termination. It is perhaps surprising then that nearly all studies of hand control tend to focus on the initial reaching and grasping component (Castiello, 2005; Castiello & Begliomini, 2008; Grafton, 2010; Turella & Lingnau, 2014), and only a ‘handful’ have examined the placement movements that often follow (Drury & Pizatella, 1983; Flindall & Gonzalez, 2019; Luabeya et al., 2024). Reach-to-grasp and reach-to-place share many similarities (e.g., eye-hand coordination, goal coding, transport kinematics, hand-object integration) but also differ in terms of intention and sensory feedback (Greenwald & Knill, 2009; Hesse & Deubel, 2010; Luabeya et al., 2024). However, at this time, it is unclear how these commonalities and differences influence the underlying cortical and subcortical networks for action.

Given the commonalities between grasp and placement, one might expect many of the neural mechanisms involved in reach-to-grasp to be relevant to placement as well. In terms of the transport component as part of the dorsomedial pathway, this might include the primary visual cortex (V1), superior parietal occipital cortex (SPOC), middle/posterior intraparietal sulcus (mIPS/pIPS), dorsal premotor cortex (PMd), primary sensory cortex (S1), and primary motor cortex (M1) (Chinellato et al., 2011; Galletti & Fattori, 2018; Karl & Whishaw, 2013; Rizzolatti & Matelli, 2003). In contrast, grasp aspects might be processed by the dorsolateral pathway and involve V1, anterior intraparietal sulcus (aIPS), ventral premotor cortex (PMv), S1, and M1 (Culham et al., 2003; Culham & Kanwisher, 2001; Vesia et al., 2017). Finally, although neuroimaging studies tend to focus on the cerebral cortex, it is well known that subcortical structures, including the cerebellum, are involved in supporting reach and grasp (Becker & Person, 2019; Izawa et al., 2012; Nowak et al., 2007; Shirinbayan et al., 2019; Thach, 2014).

The intentional differences between grasping and placing (i.e., acquiring vs. disposing of an object) might be reflected in the higher-level signals of the ‘reach’ network. In contrast, the sensory differences might evoke more specific differences in sensory regions. Indeed, grasp planning tends to be feed-forward driven, with the intention to match the hand to the object (Blohm & Crawford, 2007; Johansson & Edin, 1993), whereas placement involves different intentions (to match the object to the desired location) and additional sensory feedback at the planning and execution phases (Luabeya et al., 2026). Thus, during grasp planning and execution, vision needs to only focus on the desired object to acquire, whereas placement may require higher-level cognitive comparisons between the object and a seen or visualized target. Further, grasp starts without haptic feedback and ends with physical contact with the object, whereas place begins with haptic feedback that ends when the object is released. Each of these factors should influence cortical and subcortical sensory signals.

Further discussion of specific neural mechanisms requires speculation because direct experimental evidence is currently lacking. Neuroimaging studies have examined post-grasp actions such as object manipulation and size-weight perceptual discrepancies (Chouinard et al., 2009; Jenmalm et al., 2006), but to our knowledge, no previous imaging study has directly contrasted the mechanisms for grasping versus placing an object, either at the level of activation or at the level of functional network topology and dynamics.

To address this question, we designed an fMRI study in which participants alternated between grasping and placing a real object from / toward templates at varying locations and orientations. For a comprehensive understanding of their difference, we completed four types of analysis: 1) univariate voxelwise contrasts to established whole brain differences 2) region-of-interest (ROI) analyses based on conventional reach / grasp coordinates, 3) a multivoxel based classification analysis of our ROIs to confirm their task-dependency and 4) a graph theoretical analysis (GTA) of functional connectivity to investigate how the whole-brain distribution of signals differs between these tasks. For this analysis, we also removed global trends (the general rise in Blood-Oxygen-Level-Dependent (BOLD) activation before action), which have been suggested to reflect long-range signal correlations (Musa et al., 2025).

Given the literature and theoretical considerations provided above, we expected to find common activation across the known reach-to-grasp system, as well as local and network-level differences related to higher-level planning and sensory feedback. We found that *grasp* and *placement* share similar cortical activation patterns during the *planning* and *execution* phases, but exhibit local differences in activation, particularly in the cerebellum, that support accurate task decoding. Further, our functional connectivity analysis revealed three modules spanning the occipital, parietal, and frontal cortex, as well as the cerebellum. Overall, our results indicate that grasp and place share many similarities but differ in higher-order planning and sensory integration.

## METHODS

### Participants

Twenty-two participants passed our initial screening process in the MRI simulator suite; however, two were excluded due to discomfort in the actual MRI scanner. A total of 20 subjects (11 females, 9 males; average age, 25; SD, 4.82) participated in this study. An a priori power analysis was conducted using G*Power 3.1.9.7 (Faul et al., 2007) to determine the required sample size. Based on an effect size of 0.67 (derived from a previous task-based functional connectivity analysis; Luabeya et al., 2026), an alpha level of 0.05, and a desired power of 0.80, the analysis indicated that a total sample size of 20 participants would be required to achieve the targeted statistical significance. All participants exhibited normal or corrected-to-normal visual acuity with no reported neurological disorders and were right-handed. They provided informed consent prior to the experiment and received monetary compensation for their time. The experimental procedures were approved by the York University Human Participants Review Committee (Certificate #: e2023-336).

### Apparatus and Stimuli

An inclined platform placed on the participant’s pelvis was used to present the stimuli (Fig. 1A). The platform was positioned approximately 22 cm from the participant to facilitate a natural view and comfortable hand movements to the left and right. Participants used a black cuboid object that can be picked up and placed in a hollowed rectangular template on the platform (Fig. 1B). The template could be positioned in two locations (the top-right or the bottom-left of the platform) and could be displayed in two possible orientations: clockwise (+30°) or counterclockwise (-30°). An experimenter in the scanner wore headphones with instructions on how to change the location and orientation of the hollowed template during the intertrial interval. Two light-emitting diode (LED) lights would illuminate the target during the trial.

**Figure 1.**
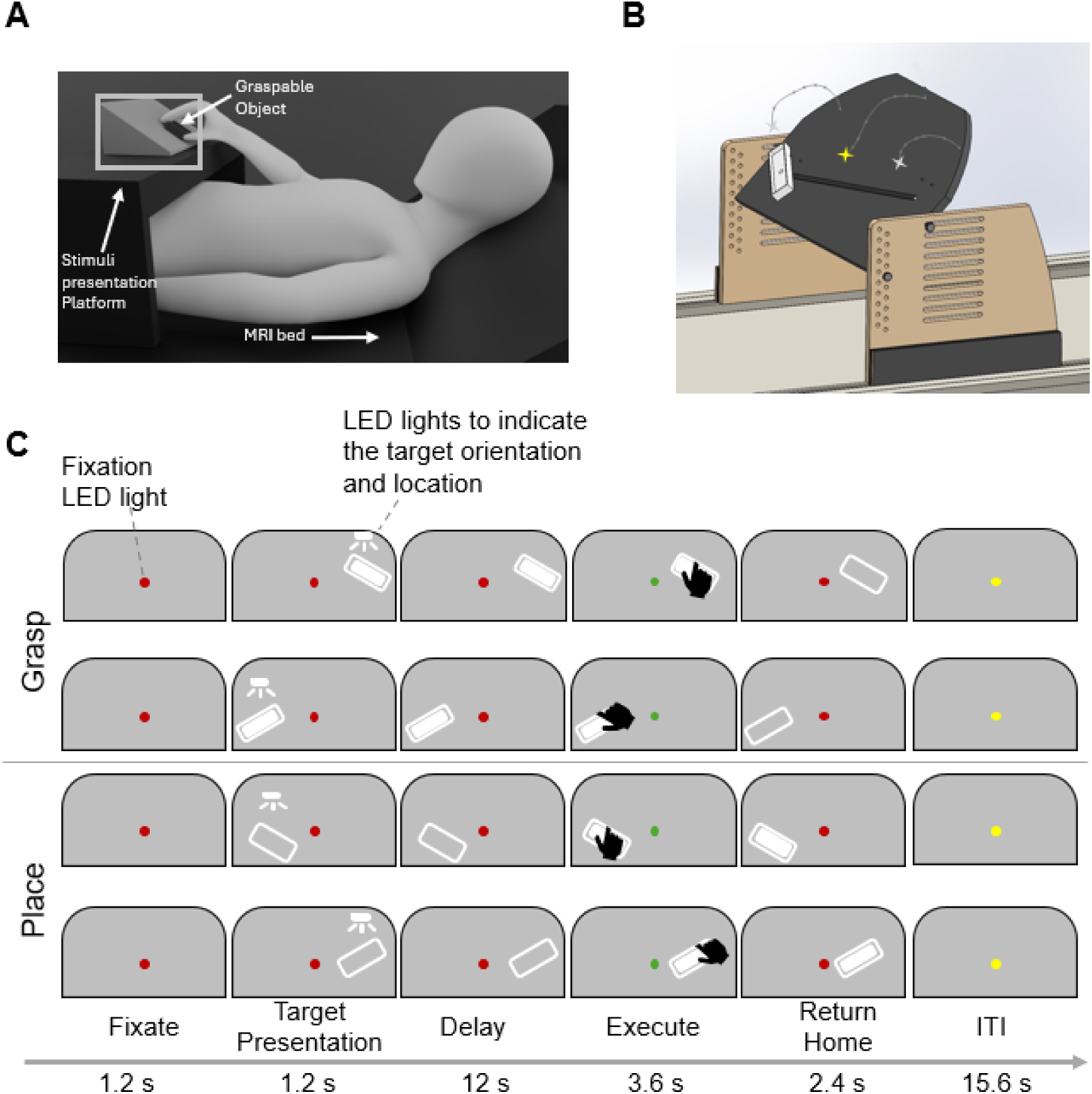
A) Experimental setup illustration (Luabeya et al., 2026). Participants were instructed to maintain fixation on the LED light at the center, and they could grasp / place the object to the left or the right. B) View of the inclined platform with the template presented to the left with the fixation light as a yellow star and the left and right LED lights as white stars. C) The time sequence shows four possible movement types: left or right *grasp* and left or right *place* with the target presented in a clockwise or counterclockwise orientation. In a trial, participants began by fixating on a red light, and a target was illuminated on the platform for 1.2 s. They waited an additional 12 s until the light turned green to perform the *grasp* or the *place*. They had 3.6 s to perform the movement and 2.4 s to return to their home position when the light turned red again. Finally, there is a 15.6 s inter-trial period before the next trial.

Participants laid in the scanner with their heads tilted (20 °) to facilitate a direct view of the objects. Foam cushions immobilized their head and upper arms to reduce motion artifacts. A belt was placed around the participant’s waist to serve as the home position, from which they started and returned after performing the movements required in the trial. To maintain similar haptic feedback during the intertrial interval, the belt featured a cuboid object that participants could hold in preparation for a *grasp* trial or rest the object on it in preparation for a *place* trial. Additionally, movement of the upper arms was restricted by a strap.

All lights on the platform and the audio to the experimenter were controlled by a custom program on a laptop PC that received signals from the MRI scanner at the start of each trial. The windows in the scanner room were blocked with black curtains, and the room’s lights remained off to improve visibility of the dimmed fixation light and the lights illuminating the target. The platform featured an infrared camera that recorded the participants’ hand movements to monitor their performance during the experiment (MRC 12M with a custom Infrared illuminator).

### Experimental Task Design

In this slow event-related fMRI task (Fig. 1B), participants grasped or placed a real 3D object to the left or right of their body midline, in clockwise or counterclockwise orientation. Participants grasped and picked up the object or placed and left it on a template on the platform. A trial consisted of three epochs: a *planning* phase after target presentation, an *executing* phase when the participant moved to place or grasp the object and return to their home position with (after *grasp* movement) or without it (after *placing* the object on the platform), and finally an intertrial period in preparation for the subsequent trial. Henceforth, the two task-relevant phases of the trial will be referred to as the *planning* and *execution*, and the two actions will be referred to as *grasp* and *place*.

Each trial began with a red fixation point, which remained visible throughout the *planning* phase. After 1.2 seconds, the target was illuminated for another 1.2 seconds, appearing on either the left or the right. This cue was followed by a 12-second delay period during which only the red fixation point was shown, allowing participants to prepare their movement without additional visual information. At the end of the *planning* phase, the fixation point turned green, serving as the go signal. Participants then had 3.6 seconds to perform either a *grasp* or a *place* movement and to hold the object at the target location until the fixation turned red again. Following this, they had 2.4 seconds to return to their home position. In the *grasp* condition, they returned with the object in hand; whereas, in the *place* condition, they returned without the object and instead held a mock block at their home position. Therefore, within a run, a *place* trial always followed a *grasp* trial, and vice versa. Each trial concluded with a 15.6-second intertrial interval, during which a yellow fixation light was displayed.

Overall, this paradigm formed a 2 (location instruction: left, right) × 2 (orientation instruction: clockwise, counterclockwise) × 2 (movement: *grasp*, *place*) design, resulting in 8 conditions. Each run consisted of 16 trials; therefore, each experimental condition was repeated twice in a pseudo-randomized order. A 14.4-second and a 13.2-second fixation point were added at the beginning and the end of each run, respectively, yielding a run time of 9.74 minutes. Each participant completed eight runs; however, for half of the participants, the odd runs began with a *place* trial, and the even runs began with a *grasp* trial, whereas the other half followed the reverse order. The sequence of conditions (*place* vs. *grasp*) was counterbalanced this way to minimize systematic bias. Additionally, an anatomical scan and a gradient echo field map were collected midway through the data collection session. Combined with setup time and a practice run, an entire session was approximately 2.5 hours.

### Imaging Parameters

Our data was collected at York University (Toronto, Canada) using a 3-T whole-body MRI system (Siemens Magnetom TIM Trio, Erlangen, Germany). To facilitate the visibility of the stimuli in the head-titled setup, the 12-channel coil of the anterior part of the 32-channel coil was replaced with a 4-channel flex coil while its 20-channel coils posterior portion was maintained (Fig. 1A).

The T2*-weighted functional data were acquired using a single-shot gradient echo-planar imaging (EPI) sequence with the following parameters: a repetition time (TR) of 1200 ms, echo time (TE) of 30 ms, a flip angle of 66°, a 240 mm field of view (FOV) and a 96 × 96 matrix, yielding a resolution of 2.5 × 2.5 × 3.0 mm³. Fifty-four axial slices were acquired with no inter-slice gap using an interleaved multi-slice mode. Slice acceleration (multiband) and GRAPPA in-plane acceleration (factor 2) were employed to accelerate acquisition. A total of 490 volumes were collected per run. A high-resolution T1-weighted anatomical image was also acquired using a 3D MPRAGE sequence (TR = 2300 ms, TE = 2.26 ms, inversion time = 900 ms, flip angle = 8°, total acquisition time = 5min 21 s) with a 256 × 256 matrix over a 256 mm FOV, yielding a 1.0 mm³ isotropic resolution across 192 sagittal slices. To correct for B0 field inhomogeneities, a gradient echo field map was acquired using a dual-echo sequence with TR = 566 ms, TE1 = 4.92 ms, TE2 = 7.38 ms, flip angle = 60°, and total acquisition time = 1min 51s with the same geometry and resolution as the functional images (2.5 × 2.5 × 3.0 mm³ voxels, 54 axial slices, FOV = 240 mm). The field map was used during preprocessing to improve spatial accuracy and reduce distortion in the EPI data.

### Preprocessing

Data were converted to NIFTI format using NITRC MRIcroGL, then they were preprocessed and analyzed using the FMRIB Software Library (FSL) (Jenkinson et al., 2012). The first three volumes of each scan were omitted to avoid T1 saturation effects. We used high-pass temporal filtering to remove low-frequency artifacts, applied spatial smoothing with a kernel size of 6.0 mm, and performed an interleaved slice scan time correction. We motion-corrected each functional run using MCFLIRT, which applied a rigid-body transformation to correct for motion artifacts (Jenkinson et al., 2002). Volumes that showed abrupt head movements over 0.9 mm were marked as outliers and regressed out (Siegel et al., 2014). Runs with 10% of volumes marked as outliers were excluded from further analysis. Functional acquisitions were coregistered to anatomical images using the Brain-Boundary Registration in the Montreal Neurological Institute (MNI152) standard space (Mazziotta et al., 1995). To confirm their compliance with the task and ensure that they are comfortable in a tight space, participants performed practice trials in an MRI simulator before the data collection session. Participants’ movements and adherence to the instructions were monitored offline. Trials were excluded from further analysis if participants did not move on time, skipped a trial, or if the experimenter’s hand was visible during the trial.

### Graph Theoretical Analysis

Since the brain is fundamentally relational, it can therefore be viewed as a network that shares information to achieve various goals and can even be considered beyond its physical connections, based on its functionality (Biswal et al., 1995; Friston, 1994). These considerations have led to the development of functional connectivity analysis and the use of mathematical approaches such as graph theoretical analysis (GTA), which represent the brain as a graph with a set of nodes (vertices) that are linked (edges) (Fig. 2 A, E) (Bullmore & Sporns, 2009; Farahani et al., 2019).

**Figure. 2:**
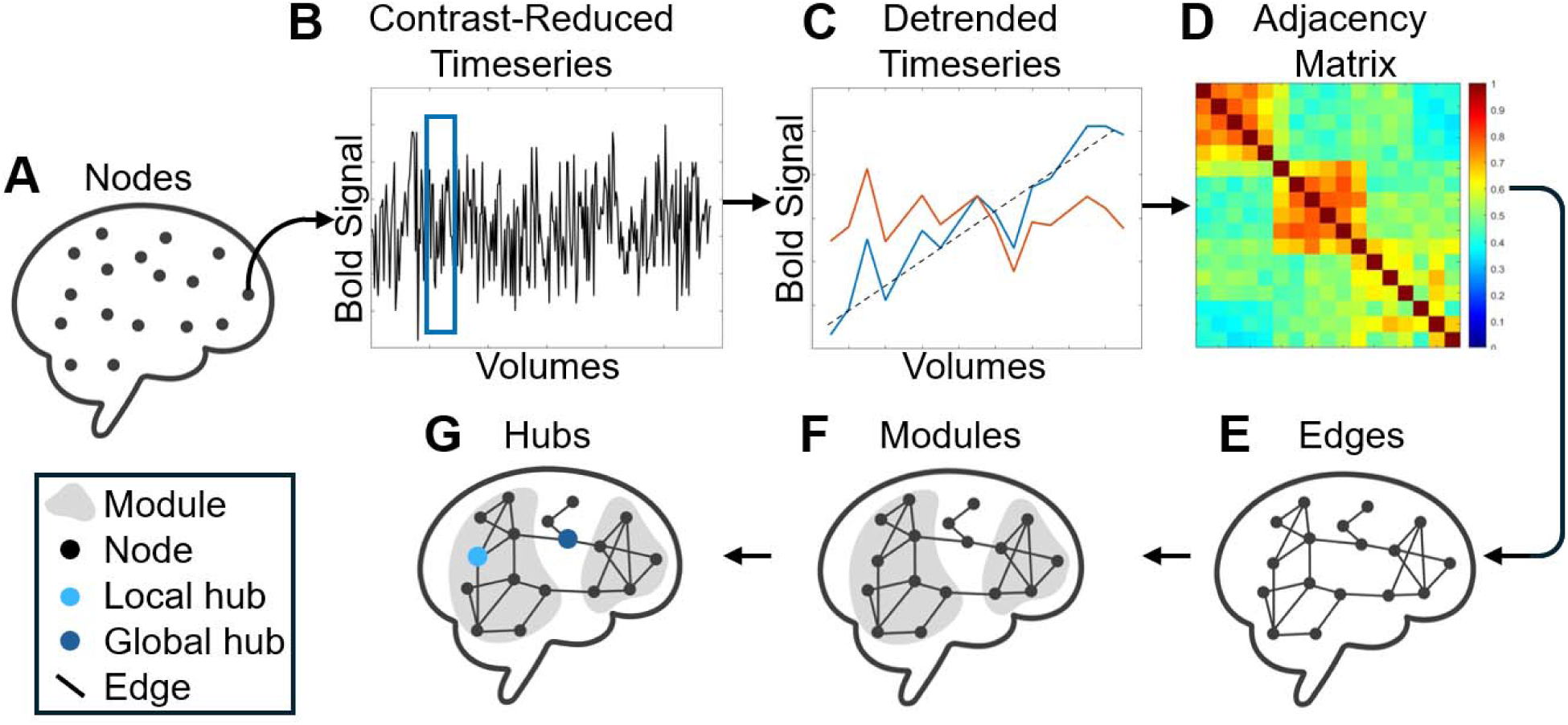
Illustration of the graph theoretical pipeline. A) Sagittal view of the brain with its set of nodes representing the different regions of interest (ROIs) from where functional data are collected. B) BOLD signal contrast-reduced timeseries is computed from an entire run for each node. Blue box corresponds to one trial extracted. C) The blue line depicts the contrast-reduced timeseries from each trial (spanning from the end of the intertrial period prior to the trial to the end of the *execution* phase), the dashed line represents its line trend, which is removed by applying a detrending algorithm. The detrended timeseries is presented in red. D) Adjacency matrix comprising the absolute value of the correlation of all the nodes in our network. These matrices are used to compute GTA measures. Blue shows nodes of low correlation, and red shows nodes of high correlation. E) The connection between the different nodes is represented by ‘edges’, the lines linking the different nodes. F) Grey areas represent the modules formed in the network. They are groups of nodes with stronger connectivity within the modules than outside the modules. G) Represents nodes of particular importance. There are eigenvector centrality hubs, which have a node with a strong connection to other highly connected nodes in blue, and betweenness centrality hubs, which serve as bridges connecting different clusters of nodes in red.

To achieve this graph representation of our network, we create 6mm diameter spheres on FSLeyes for our 200 cortical regions of interest (ROIs) and 32 cerebellar ROIs. These coordinates were identified using Yeo’s 17-network atlas with a 200-node cortical parcellation and a 32-node cerebellar parcellation (Nettekoven et al., 2024; Schaefer et al., 2018). Considering that both cortical and subcortical regions provide a comprehensive assessment of their implications in motor control, as the cerebellum is important for timing, prediction, and coordination (Bastian, 2006; Salman, 2002; Sokolov et al., 2017). The raw BOLD signals from these 232 spheres, for each run across all participants included in our analysis, were extracted to conduct graph-theoretical functional connectivity analysis (Fig. 2B). For these runs, we also obtained the event files, parameter estimates, and residuals for each region of interest to compute the contrast-reduced time series of our data. This approach was taken to remove elements irrelevant to our task (for example, the color of the fixation point), which might otherwise obscure our findings (Musa et al., 2025; Luabeya et al., 2026). Because we intended to understand the difference between grasp and place, we isolated 17 volumes representing our events of interest for each trial in a run (Fig. 2C). During a trial, our 17 datapoints included the two volumes before the start of the trial and ran until the initial movement to the platform to grasp or place the object. Then, we correlated the extracted contrast-reduced timeseries of the nodes against each other for all the trials (Fig. 2D). The correlation outputs yielded 232 x 232 adjacency matrices. We computed their absolute values to confirm that we are limiting our analysis to the amplitude of the correlations, not their directionality (Ran et al., 2020). We calculated the grand average across all trials to create one undirected weighted adjacency matrix per condition. Each value in the matrix represents the strength of the functional connection ‘edge’ between two nodes (Fig. 2E).

Network analysis using GTA can be understood at three different levels: 1) a macro/global scale, which views the entire network as one conglomerate system, 2) a meso/intermediate scale, which looks at the subnetwork that forms communities in the system, 3) a micro/local scale, which identifies the individual nodes leading the communication in the network. At the global level, the *clustering coefficient* assesses the network’s ability to facilitate the formation of highly connected subnetwork cliques to optimize information sharing; the *global efficiency* quantifies how effectively the network exchanges information; and the *energy* estimates the stability of network synchronization (Demaine et al., 2006; Ghaderi et al., 2020; Latora & Marchiori, 2001; Rubinov & Sporns, 2010). Overall, these measures allow one to evaluate the integration and segregation of the network as a whole unit. At the meso-scale, to maximize communication efficiency and reduce the effort required to share information, the brain uses a small-world organization, where nodes are grouped into smaller communities with greater communication (Newman, 2006; Sporns & Betzel, 2016). The modularity analysis allows the identification of these communities (modules) with high connectivity within the subnetwork rather than across subnetworks (Fig. 2F). These communities reveal how our network segregates functionally to achieve a more effective information flow (Latora & Marchiori, 2001). At the local scale, to identify which nodes are particularly important for communication within and between modules, we computed two centrality measures (eigenvector centrality and betweenness centrality; Fig 2G). For this purpose, and to quantify the quality of our module, we combined the trial-by-trial adjacency matrices for each participant, yielding a unique averaged adjacency matrix for each condition. We then computed eigenvector centrality to identify local hubs within modules and betweenness centrality to identify global hubs driving communication across modules (Bonacich, 2007; Freeman, 1977).

Then, centrality measurements were computed on these matrices using the Brain Connectivity Toolbox (Rubinov & Sporns, 2010). We assess the modularity score of each module by calculating Newman’s Modularity (Q) using an open-access ‘MODU’ function (Ghaderi et al., 2020, 2025; Newman, 2006).

Sensorimotor studies have often reported a “buildup” or ramping of BOLD activation over time during a trial. Musa and colleagues (2025) have shown that detrending (removing the linear trend present in the data) reduces long-range communication and focuses more on local processes (Fig. 2C). We wanted to use a similar approach to investigate how the formation of modules might be impacted by the absence of ‘global communication’. Therefore, we also computed the modularity analysis in this detrended time series.

### Statistical Analyses

#### Univariate Analysis

Data analyses were performed using the general linear model conducted on FSL FEAT (version 6.00). Each run was modeled using a voxelwise GLM, with each predictor derived from a boxcar wave function convolved with FSL’s canonical double-gamma hemodynamic response function, and temporal derivatives included to account for variability in response timing. For runs that were not completely excluded from further analysis but contained behavioral errors (e.g., hand movement during the *planning* phase), we included a predictor to regress out these trials (corresponding to 36 trials in total). We modeled our predictors using these two time periods (*planning* and *execution* phases) based on movement type (*grasp* or *place*), resulting in a total of four predictors of interest: Planning-Grasp, Planning-Place, Execution-Grasp, and Execution-Place. We then used a fixed-effect model to combine the valid runs within subjects. Finally, we used a non-parametric permutation test with 5000 permutations to conduct group analysis. We applied FSL randomise with a Threshold-Free Cluster Enhanced (TFCE) approach to correct for multiple comparisons and produced a Family-Wise Error corrected p-value < 0.05 (Worsley, 2001). We compared them to the intertrial interval, which served as the baseline for identifying brain areas that showed increased activation during the *planning* and *execution* phases. After performing the permutation test for our two conditions to specify areas with conjoined increases in activity, we identified the minimum significant cluster size and used the largest cluster size to define significant join activation in their intersection map for *planning* and *execution* phases. We also contrasted *grasp* and *place* during the two phases to identify regions that showed increased activity specific to the movement type.

#### Regions-of-Interest and Classification Analyses

Region-of-interest (ROI) analysis was conducted across seven action-related areas in the left hemisphere, as all movements were performed with the right hand, using RStudio 2024.12.1. These regions included the primary visual cortex (V1), posterior intraparietal cortex (pIPS), anterior intraparietal cortex (aIPS), primary sensory cortex (S1), primary motor cortex (M1), ventral premotor cortex (PMv), and dorsal premotor cortex (PMd). Their coordinates were identified from the neurosynth.org meta-analysis using the keywords grasping and reaching, and V1 coordinate was defined from previous grasping work (Monaco et al., 2020), which were converted to MNI space using seed-based d mapping smproject.com according to Lancaster and colleagues (2007). Their values were averaged over a 6mm radius sphere centered on the centroid coordinates. We extracted parameter estimates for the four predictors (Planning Grasp, Planning Place, Execution Grasp, and Execution Place) across the 7 ROIs to assess their ability to dissociate between grasp and place in a classification analysis. We conducted t-tests to compare the beta weight in our two conditions (*grasp* vs *place*) and used false discovery rate (FDR: q) to correct for multiple comparisons.

Afterward, we attempt to identify how well these regions can dissociate between our two main conditions. We conducted trial-by-trial extraction of the beta weight for each ROI after z-normalizing its 3D image to remove run-specific amplitude differences. We removed outliers relative to the mean, resulting in a total of 122 voxels per ROI per trial to feed the same number of voxels to our classifier. We applied them in a supervised classification analysis using MATLAB’s Classifier Learner app, which evaluates different algorithms to identify the most efficient approach for distinguishing between our two conditions. To confirm that the trained classifier could generalize to an untrained dataset, we used 10-fold cross-validation (Valente et al., 2021; Varoquaux et al., 2017; Weaverdyck et al., 2020). This machine learning approach is a robust resampling technique where the data is divided into 10 parts (’fold’), then trained on 9 folds and tested on the last one. The step is repeated in 10 iterations so that each fold is tested at least once. Then, we used Kruskal-Wallis algorithm to rank the importance of each feature introduced to the model in their classification effort. Therefore, providing a way to identify which ROIs and phases carried meaningful information to represent the movement type.

#### GTA Statistics

To understand how grasp and place information is shared functionally, we use modularity function in the BCT toolbox at the standard resolution of 1 and assign each node to a module. To statistically confirm that subnetworks are nonrandom modules whose modularity exceeds that expected under a null model, we created a randomized subnetwork of the same size. We then computed the mean log-transformed of the normalized modularity score and applied it to a one-sample sign-flip permutation test (right-tailed with 50000 permutations). To compare our two conditions, we also conducted a nonparametric permutation t-test with 10000 randomizations for our modularity score and corrected for multiple comparisons using a false discovery rate (FDR: q) (Tomou et al, Luabeya 2026). We calculated our centrality measures for the unmodified *grasp* and *place* conditions to determine which nodes serve as hubs that lead intra- (eigenvector centrality) and inter-module (betweenness centrality) communication. To identify reliable hubs and avoid circular analysis, the centrality data were split into two groups. The first subset selected candidate hubs, defined as nodes with centrality measures at or above the 95th percentile. The second subset was tested for significance by conducting a one-sample sign-flip permutation test, comparing the real centrality dataset to null models and generating centrality scores. To increase stability in our analysis, we applied split-half validation with 100 iterations and defined hubs as nodes that were reliably selected as part of the 95th percentile and identified as statistically significant compared to null in at least 60 of the 100 validation iterations.

## RESULTS

The aim of this study was to compare and contrast brain activity and functional connectivity during *grasping* versus *placing* movements. We first performed a voxelwise univariate analysis to identify brain regions that showed overlapping or different activation between conditions at the whole-brain level. Next, we conducted a region-of-interest (ROI) analysis to focus on specific cortical areas previously implicated in object-directed actions, providing more targeted insights into task-related activity patterns. We then applied a decoding classifier to determine whether the observed activity patterns could reliably distinguish between *grasp* and *place* movements, providing a data-driven validation of task-specific neural signatures. Finally, we performed a GTA functional connectivity analysis to investigate how the brain organizes itself functionally to execute our two action types, using both original and detrended datasets (Fig. 2C). All brain networks were visualized using BrainNet Viewer (Xia et al., 2013).

### Univariate Analysis

#### General Activation during Planning Phase

To identify general *planning* activity across our two tasks, we computed increases in BOLD activation relative to baseline for *grasp* (Fig. 3 A), for *place* (Fig. 3 B) and their conjunction (Fig. 3 C) with P_FWE_ < 0.05. We observed a widespread bilateral increase in activity for both *Grasp* and *Place* tasks, as confirmed in the conjunction analysis (Fig. 3 C). This included Prefrontal Cortex (PFC), Superior Frontal Gyrus (SFG), insula, cingulate cortex, Supplementary Motor Area (SMA), Superior Temporal Gyrus (STG), Inferior Parietal Lobule (IPL), Middle Temporal Gyrus (MTG), and Early Visual cortex (EVC). However, left PMd, M1 and S1 showed a significant increase in activity for *Place* that was not observed in *Grasp* or conjunction analysis. During *planning,* the cerebellar vermis (VI-X), the left cerebellar lobules (I-VI, VIIb, IX, Crus I, Crus II), showed increased activation for both *Grasp* and *Place*; whereas the right cerebellum showed a smaller overlapping activation in the right lobules (VI and Crus I). Overall, our two conditions shared many similarities across the cortex and cerebellum, but also hinted at differences that we will focus on further below.

**Figure 3.**
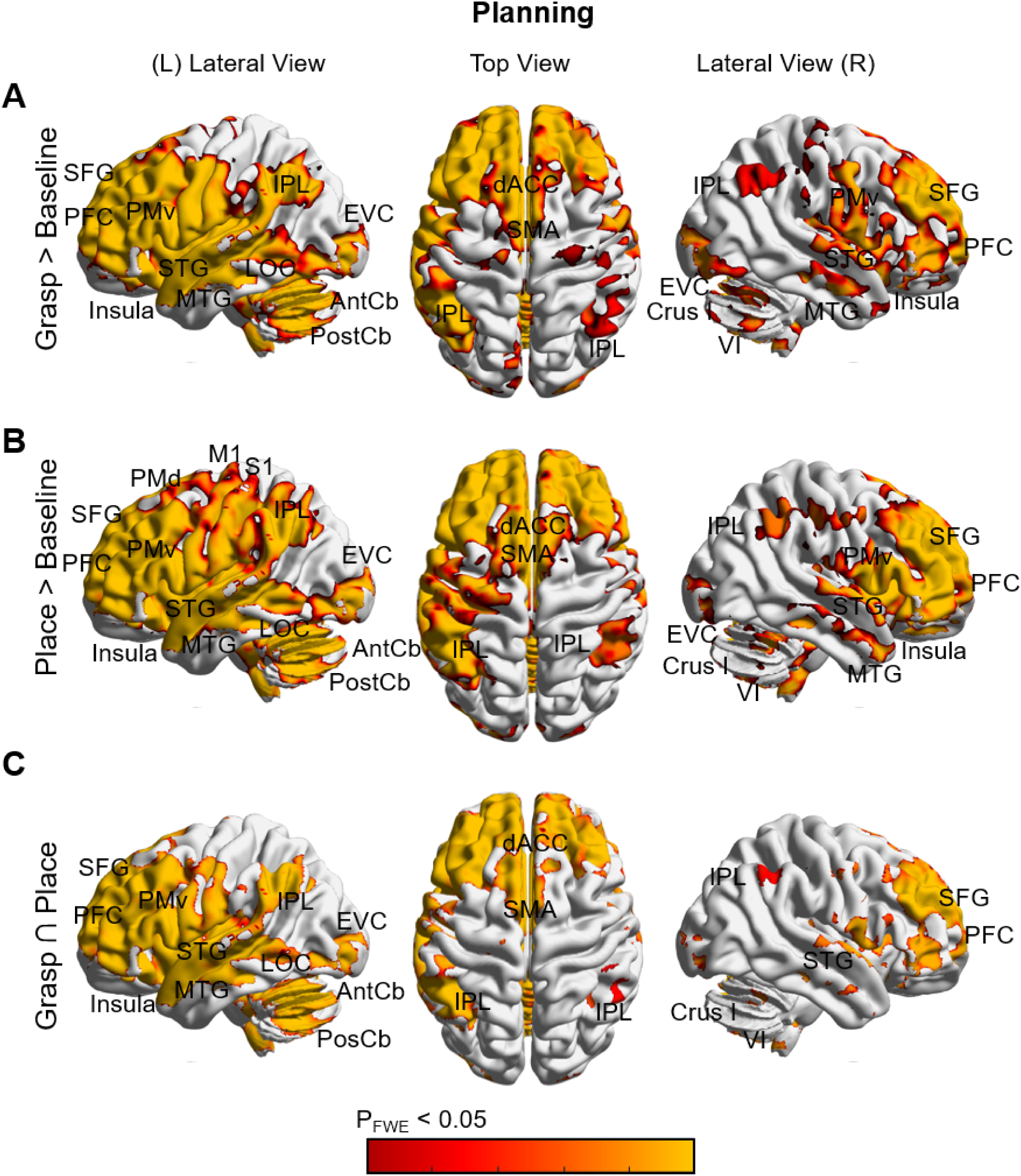
Cortical activation maps for the *Grasp* (A), *Place* (B) conditions and their conjunction (C) during the *planning* phase. Color bars indicating statistical intensity from a non-parametric permutation test with 5000 permutations (with a P_FWE_ < 0.05). Warmer colors indicate greater activation than baseline. Lateral and medial views of both hemispheres are shown for each condition. Abbreviations: dACC: dorsal anterior cingulate cortex, EVC: Early visual cortex, LOC: Lateral occipital cortex, M1: Primary motor cortex, MTG: Middle temporal gyrus, PFC: Prefrontal cortex, PMd: Dorsal premotor cortex, PMv: Ventral premotor cortex, IPL: Inferior parietal lobule, S1: Primary somatosensory cortex, SFG: Superior frontal gyrus, SMA: Supplementary motor area, SPOC: Superior parieto-occipital cortex, STG: Superior temporal gyrus, AntCb: Anterior cerebellar lobe, PostCb: Posterior cerebellar lobe.

#### General Activation during Execution Phase

Again, we computed increases in activity relative to baseline for *Grasp* (Fig. 4 A), *Place* (Fig. 4 B) and their conjunction (Fig. 4 C) (for P_FWE_ < 0.05) to identify similarities and differences during *execution* phase. We observed a widespread bilateral increase in activity for *grasp* and *place* (confirmed in the conjunction) in the S1, PMd, PMv, SMA, Superior Parietal Lobule (SPL), SPOC, LOC, insula, and cingulate cortex. In addition, we found lateralization with greater activity in left M1 (i.e., opposite hemisphere to the hand used). In the cerebellum, there was a wide bilateral activation in the anterior lobules (I-V), the posterior lobules (VI-X, Crus I and II) for both *grasp* and *place*. Overall, outside of the sensorimotor cortex, activation was more lateralized during the *planning* phase and bilateral during the *execution*. Furthermore, there was less frontal activation but more sensorimotor activation during the *execution* phase.

**Figure 4.**
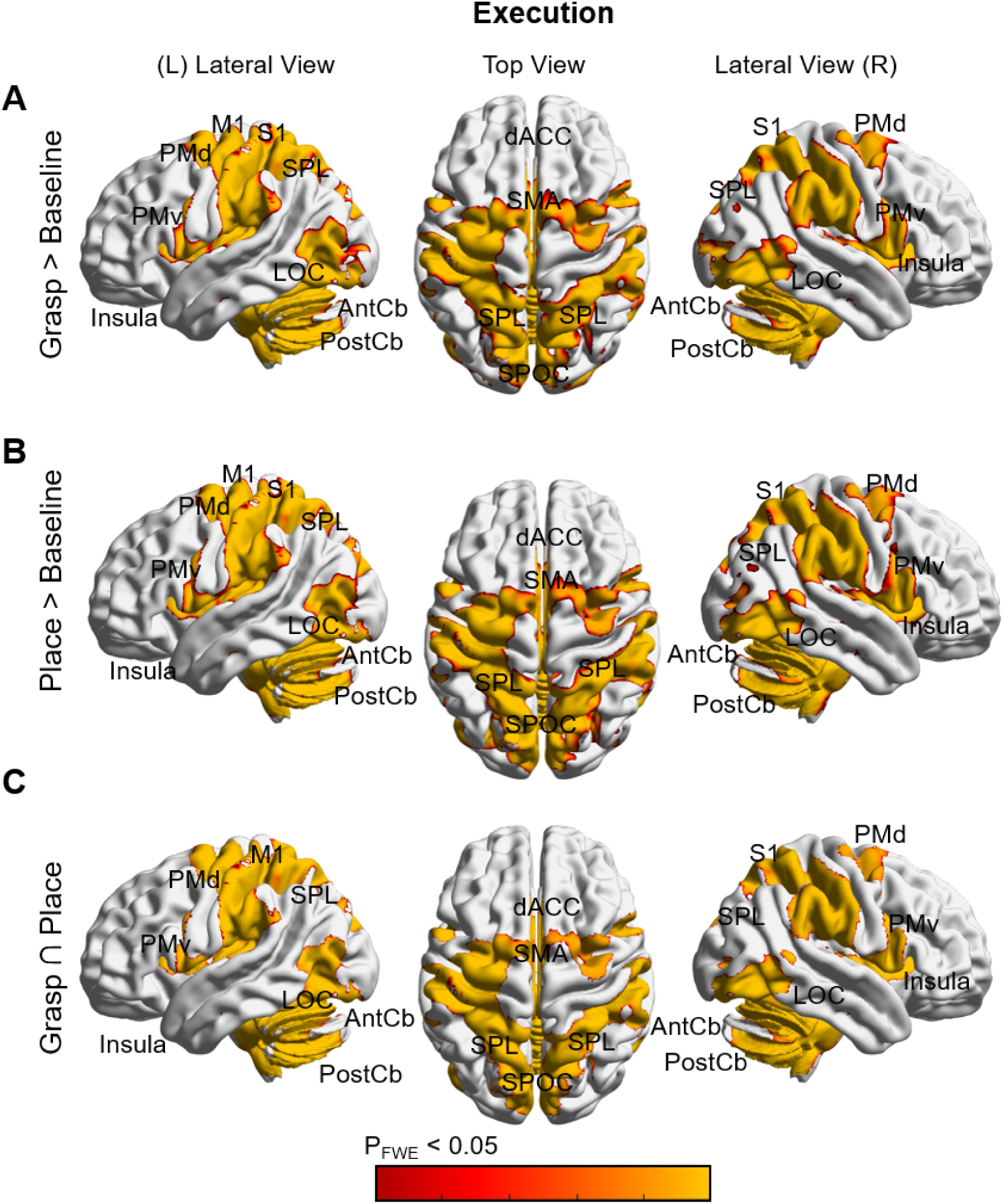
Cortical activation maps for the Grasp (A), *Place* (B) conditions and their conjunction (C) during the *execution* phase. Color bars indicating statistical intensity from a non-parametric permutation test with 5000 permutations (with a P_FWE_ < 0.05). Warmer colors indicate greater activation than baseline. Lateral and medial views of both hemispheres are shown for each condition. Abbreviations: dACC: dorsal anterior cingulate cortex, EVC: Early visual cortex, LOC: Lateral occipital cortex, M1: Primary motor cortex, MTG: Middle temporal gyrus, PMd: Dorsal premotor cortex, PMv: Ventral premotor cortex, SPL: Superior parietal lobule, S1: Primary somatosensory cortex, SMA: Supplementary motor area, SPOC: Superior parieto-occipital cortex, AntCb: Anterior cerebellar lobe, PostCb: Posterior cerebellar lobe.

#### Differences Between Reach and Place Activity

To directly distinguish task-specific activity, we performed a Grasp vs Place contrast across the two task epochs. During the *planning* phase, there was a significant increase in the left lingual gyrus for *grasp* condition, and two cerebellar locations for *place*: Right lobule VIIIa and the Right lobule V (Fig. 5A). During the *execution* phase, there were task-specific increases in activation for *placement* in the left and right occipital cuneus, occipital fusiform gyrus and lingual gyrus (Fig. 5B). The cerebellum showed bilateral *place*-related preference in the spinocerebellum (lobules I-X and Crus II) and the cerebrocerebellum (lobules I-IX). These findings suggest that grasp and place exhibit regional specialization, which we explore further in the next section.

**Figure 5.**
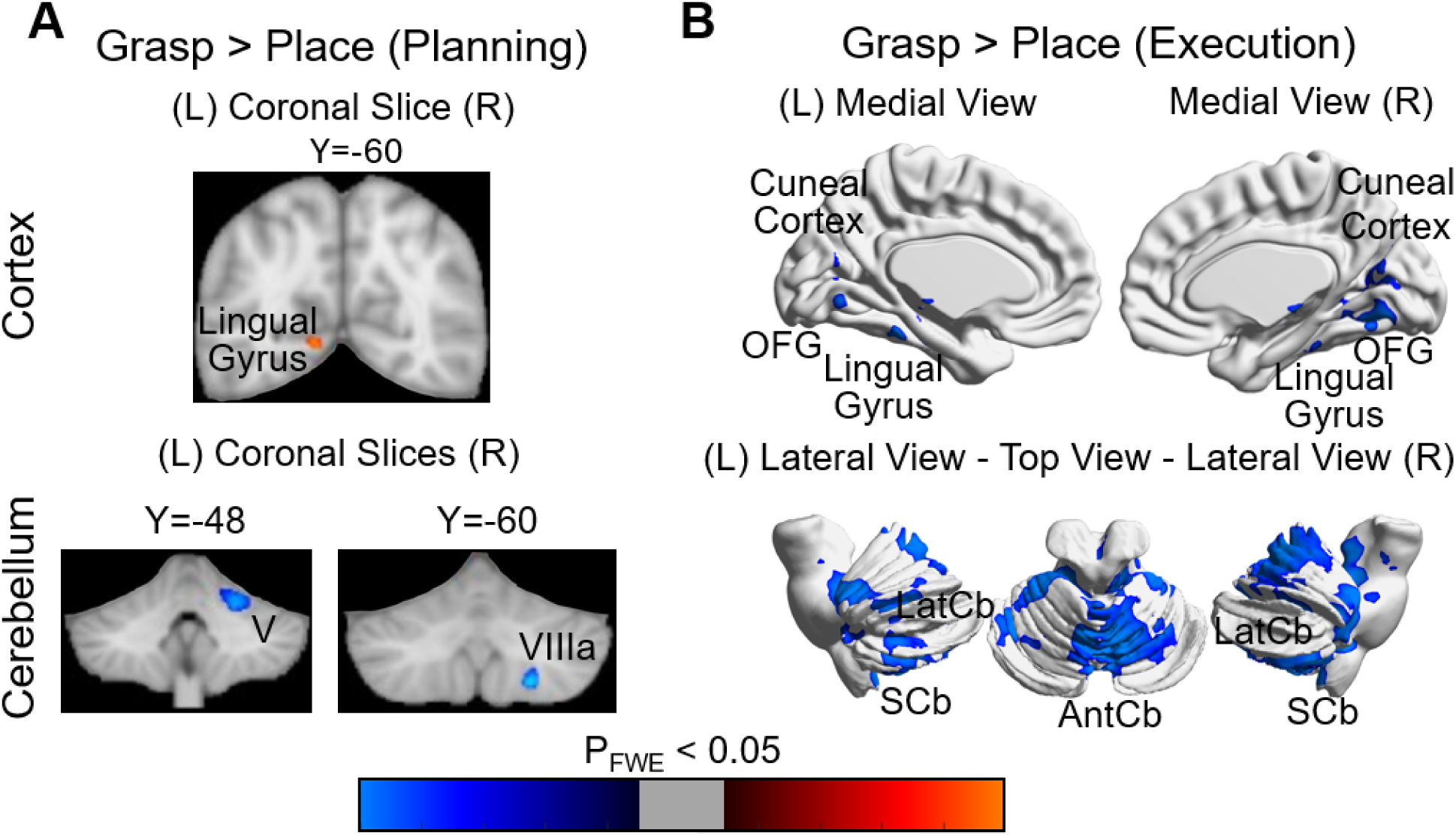
Cortical activation maps for the *Grasp* vs *Place* comparison during the *planning* phase (A) and the *execution* phase (B). Color bars indicating statistical intensity from a non-parametric permutation test with 5000 permutations (with a P_FWE_ < 0.05). Cool colors represent a significantly greater activation for *Place*, and warm colors depict a significantly greater activation for *Grasp*. The top row shows cortical activation from a coronal slice (Y_MNI_ = -60) during *planning* and the bilateral medial views during *execution*. The bottom row shows cerebellar activation from two coronal slices (Y_MNI_ = - 48 and Y_MNI_ = -60) during *planning* and the lateral and top views during the *execution*. Abbreviations: OFG: Occipital Fusiform Gyrus, AntCb: Anterior lobe of cerebellum, V: Cerebellar Lobule V, VIIIa: Cerebellar Lobule VIIIa, LatCb: Lateral cerebellum (cerebrocerebellum), SCb: Spinocerebellum.

### Region of Interest Analysis

To examine task-related brain activity within known action-related areas, we conducted paired-sample t-tests on β weights extracted from 7 ROIs in the left hemispheres to compare *grasp* and *place*. During the *planning* phase (Fig. 6A), *placement* showed a significant increase compared to *grasp* in aIPS (mean difference = 0.071, t(19) = 2.400, q = 0.026), S1 (mean difference = 0.044, t(19) = 3.129, q = 0.005), M1 (mean difference = 0.049, t = 2.427, q = 0.025) and PMd (mean difference = 0.077, t(19) = 2.749, q = 0.013) compare to *grasp*, but not in V1 (mean difference = 0.028, t(19) = 0.854, q = 0.434), pIPS (mean difference = 0.013, t(19) = 0.925, q = 0.434) and PMv (mean difference = 0.015, t(19) = 0.798, q = 0.434). None of the seven ROIs showed significant task modulations during the *execution* phase (Fig. 6B) after false discovery rate (q > 0.05).

**Figure 6.**
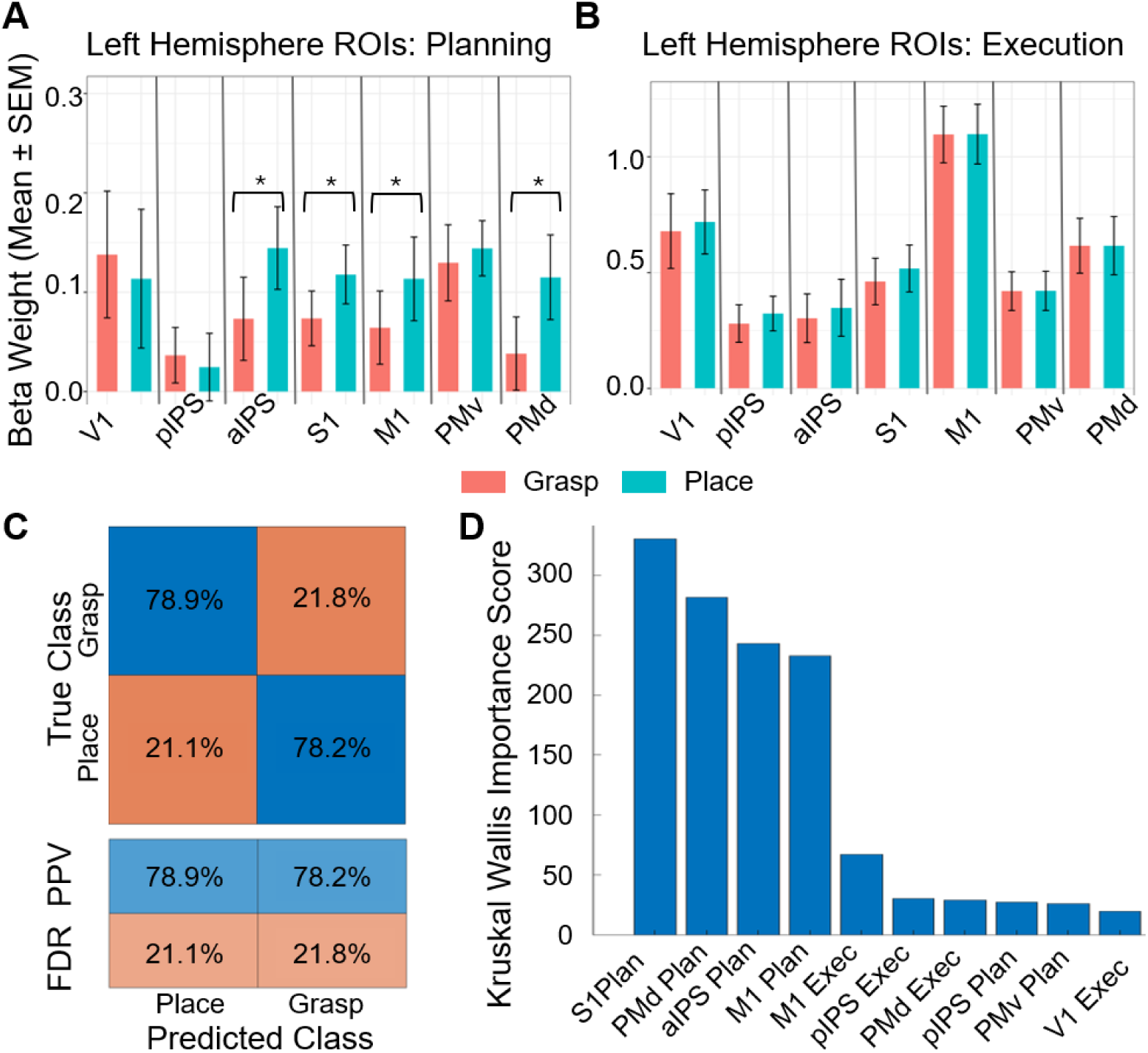
Bar graphs of the beta weights from 8 ROIs in the left hemisphere during the *planning* phase (A) and the *execution* phase (B). Asterisks show the significant difference between *grasp* and *place* at p < 0.05 after FDR correction. C) Multivoxel Classifier Analysis represents the performance of the k-Nearest Neighbor algorithm to differentiate between *Grasp* and *Place*. The top table panel displays the confusion matrix, comparing the predicted data point identity (*grasp* or *place*) with its true identity. The bottom panel table shows the positive predictive values (PPV) and false discovery rates (FDR) of the *Grasp* data (left column) and the *Place* data (right column). D) The Kruskal-Wallis importance scores of the ROIs and time period (Plan = *planning* phase, Exec = *execution* phase) to classify between *grasp* and *place*.

### Classification Analysis

We used classifier models to see whether these action-related regions of interest (Fig. 5a) could differentiate between *grasp* and *place*. Among the models tested, the weighted K-Nearest Neighbors (KNN) achieved the highest accuracy of 78.53% in distinguishing between *grasp* and *place* during 10-fold cross-validation. KNN is a supervised algorithm that classifies based on its proximal neighbors, placing greater emphasis (weight) on the closest neighbors (Ghaderi et al., 2023a). Outperforming the traditional Support Vector Machine (SVM) algorithm, as weighted KNN is less affected by outliers, and is more suitable for complex, flexible data. The classification performance was summarized in our confusion matrix, which shows the distribution of true and predicted class labels (Fig. 6C). The true class reflects class-specific sensitivity, and the predicted class reports the positive predictive value (PPV) and the false discovery rate (FDR). The model achieved a PPV of 78.9% (95% Confidence interval (CI) [78.3 – 79.5]) with an FDR of 21.1% (CI [20.5 – 21.7]) for the *Grasp* movement, and a PPV of 78.25% (CI [ 77.6 – 78.8]) with an FDR of 21.8% (CI [21.3 – 22.4]) for the *Place* movement. The Kruskal-Wallis (KW) feature selection analysis revealed that the information obtained during *planning* phase from S1 (KW score = 330.67), PMd (KW score = 281.70), aIPS (KW score = 243.10), and M1 (KW score = 232.68) ranked as the most informative in order to distinguish between *grasp* and *place* at an Importance Score greater than 100 (Fig. 6D).

### Graph Theory Analysis

To compare functional connectivity in our two tasks at the whole brain level, we used a graph-theoretical analysis (GTA) approach and correlated the contrast-reduced time series of our 232 cortico-cerebellar nodes (Fig. 2). We did not find any significant difference between commonly used global parameters (Clustering Coefficient, Global Efficiency and Energy) for the reach vs. placement networks (Supplementary Fig. 1), so instead focus on the mesoscale and local aspects of functional network organization. We also examined the influence of detrending the data (Fig. 2C), which has been shown to reduce large-scale signal correlations (Musa et al., 2025).

#### Modularity Analysis: Original Data

To understand the mesoscale properties of these networks, we conducted a modularity analysis, which grouped our nodes into modules/subnetworks, with higher connectivity within modules than across modules (Fig. 7A-B). We identified three modules in the original *grasp* and *place* conditions: a Cerebello-Occipito-Parietal Module (Module 1: red), a Prefrontal Module (Module 2: blue) and a Sensorimotor Module (Module 3: purple). For the original (unmodified) timeseries, the node assignments were identical between *grasp* and *place*.

**Figure 7.**
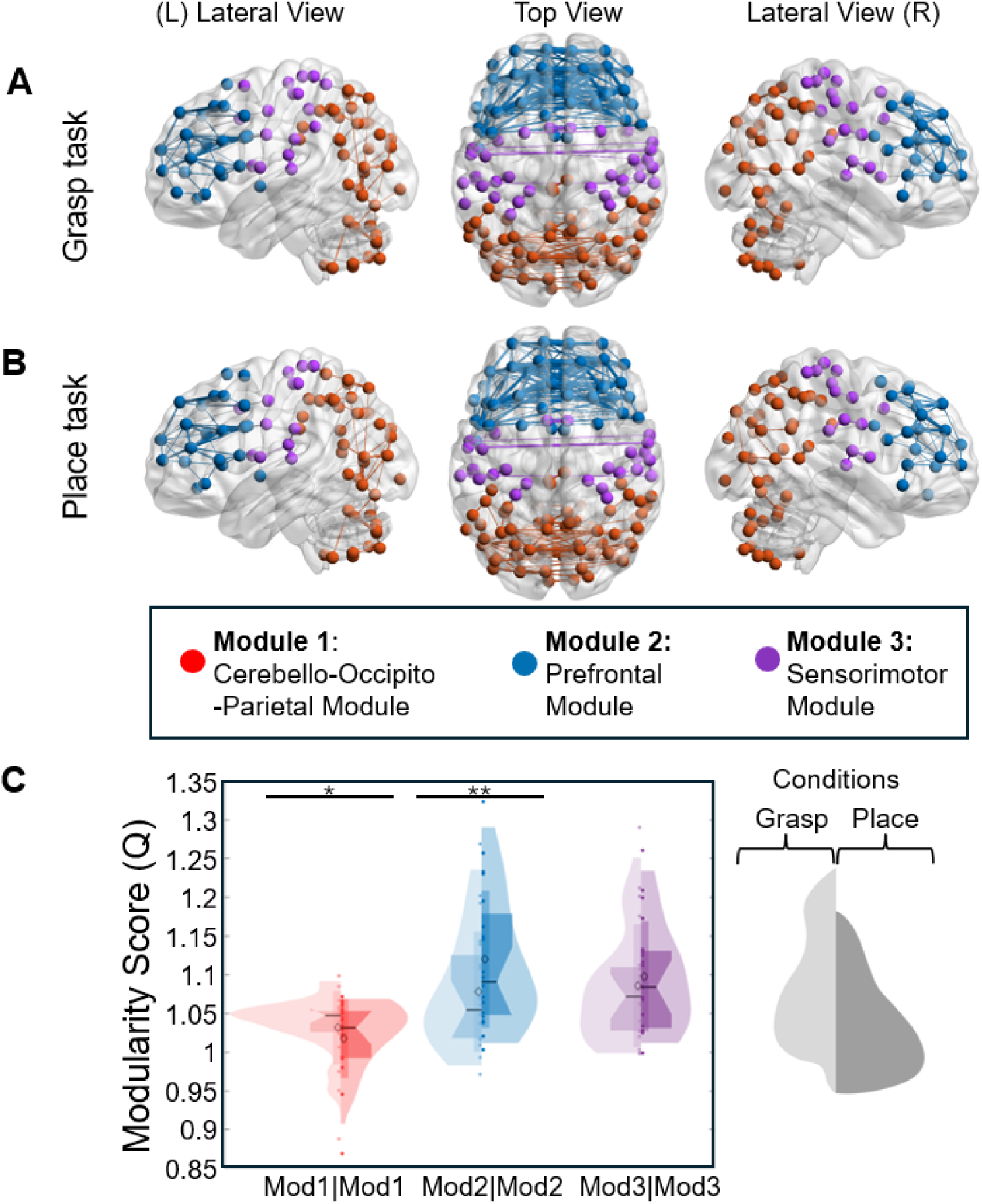
Trend unmodified network modularity analysis. A) represents the three module groupings for the *grasp* task. B) represents the three module groupings for the *place* task. C) Violin plots of the Modularity score of each module from the networks. The *Grasp*’s modules are presented in the lighter colors to the left and the *Place*’s modules are presented in the darker colors to the right (Mod = Module). Modules are color-coded: Cerebello-Occipito-Parietal Module (Module 1) in red, the Prefrontal Module (Module 2) in blue and Sensorimotor (Module 3) in purple. The FDR-corrected significant differences between conditions are indicated by asterisks with *: p < 0.05, **: p < 0.01, and ***: p < 0.001.

We then used the modularity score to quantify how strong the communities were compared to a random subnetwork of the same size and whether their community strengths differed depending on the task executed (Fig. 7C). We found that all three modules performed better than a null model for the *grasp* condition (Module 1 q = 0.0062, Module 2 q = 5.90E-5, Module 3 q = 5.90E-5). However, only two of the three modules had scores significantly greater than the null model for the *place* condition (Module 1: q = 0.0866; Module 2: q = 2.99E-5; Module 3: q = 2.99E-5).

When the modules are compared based on condition (Fig. 7C), we found that Module 1 and Module 2 showed a significant difference, with *grasp* having a higher modularity score than *place* for Module 1 (t(19) = 2.7395, q = 0.0139) and inversely, *place* having a higher modularity score than *grasp* in Module 2 (t(19) = -3.6379, q = 0.0045). Whereas Module 3 did not show a significant difference in modularity score (t(19) = -0.9754, q = 0.3447). Overall, the data suggest that *grasp* and *place* are grouped into similar three ‘real’ nonrandom modules in the network, where the sensorimotor module was task-independent, the Cerebello-Occipito-Parietal module showed a higher modularity score for *grasp*, and the Prefrontal module showed a higher score for *placement*.

#### Modularity Analysis: Detrended Data

To assess how the general increase in BOLD response influences modularity, we computed the modularity score in datasets with detrended contrast-reduced timeseries. After detrending, the nodes were grouped into three modules for the *grasp* condition (Fig. 8A), similar to those for the original dataset. When tested against a null model, their modularity scores remained significantly higher than chance (Module 1 q = 0.0069, Module 2 q = 1.20E-4, Module 3 q = 5.99E-5).

**Figure 8.**
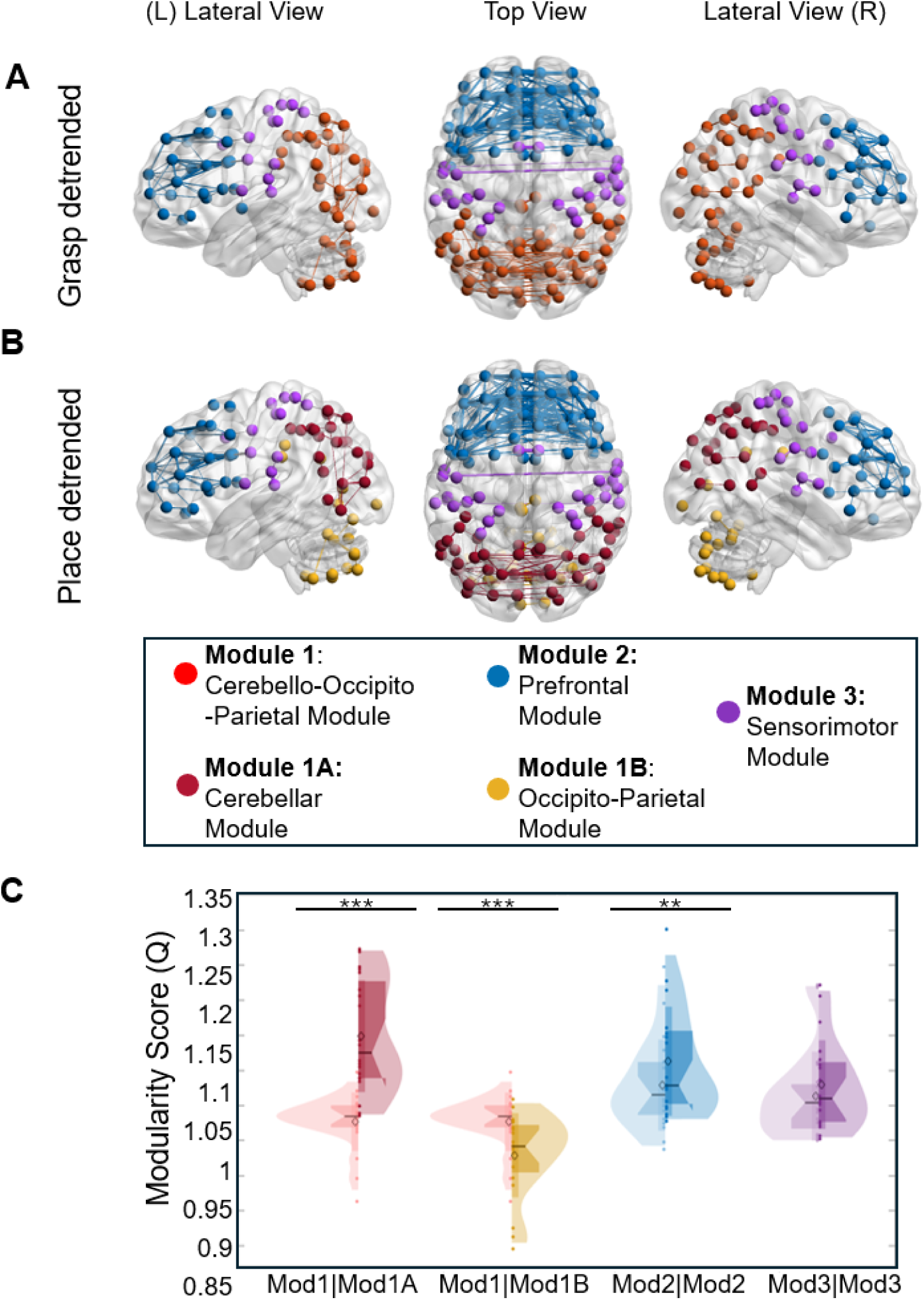
Detrended Network modularity analysis for the *Grasp* and *Place* conditions. A) represents the three module groupings for the *grasp* task. B) represents the four module groupings for the place *grasp* task. C) Violin plots of the Modularity score of each module from the networks. The *Grasp*’s modules are presented in the lighter colors to the left and the *Place*’s modules are presented in the darker colors to the right (Mod = Module). Modules are color-coded: Cerebello-Occipito-Parietal Module (Module 1) in red, Prefrontal Module (Module 2) in blue and Sensorimotor (Module 3) in purple for the *grasp* side, and Cerebellum Module (Module 1A) in burgundy, Occipito-Parietal Module (Module 1B) in yellow, Prefrontal Module (Module 2) in blue and Sensorimotor (Module 3) in purple for the *place* side. The FDR-corrected significant differences between conditions are indicated by asterisks with *: p < 0.05, **: p < 0.01, and ***: p < 0.001.

In contrast, the nodes in the *place* condition were grouped into four modules (Fig. 8B): an Occipito-Parietal Module (Module 1A in dark red), a Cerebellar Module (Module 1B in yellow), a Prefrontal Module (Module 2 in blue), and a Sensorimotor Module (Module 3 in red). When tested against a null model, only three of these modules were considered to be nonrandom (Module 1A q = 2.666E-5, Module 1b q = 0.9625, Module 2 q = 2.666E-5, Module 3 q = 2.666E-5). These data suggest that, after detrending, the network’s integrative behavior decreases in the *place* condition relative to the *grasp* condition.

To quantify these differences, we again computed modularity scores (Fig. 8C). Compared to Module 1 (*Grasp*) Module 1A (*Place*) showed a significant increase in modularity for the *place* condition (t(19) = -6.7144, q = 1.998E-4), whereas, compared with Module 1b showed a lower modularity score (t(19) = 5.294, q = 1.998E-4). Module 2 still showed a significant increase in its modularity strength for *place* in comparison to *grasp* (t(19) = -3.4168, q = 0.0027), and Module 3 did not have a significant difference between *grasp* and *place* (t(19) = -1.387, q = 0.1871). These results suggest that removing the global trend amplifies the differences between the *grasp* and *place* networks.

#### Network Hubs: Local vs. Global Connectivity

We used two centrality measures (eigenvector and betweenness) to identify hubs: nodes that are highly correlated with other highly correlated nodes, and nodes that served as important bridges connecting other pairs of nodes, respectively. To be qualified as hubs, our nodes had to 1) fall within the top 95^th^ percentile of centrality measures among all nodes, and 2) show statistically significant higher centrality compared to the corresponding node in a null model. Therefore, the hubs are not derived from a direct comparison between our conditions but by contrasting them with randomized network configurations.

*Eigenvector centrality* was used to identify nodes that were highly correlated within a module (Fig. 9A-B). In the Grasp condition (Fig. 9A), we identified eight eigenvector hubs, with three hubs located in the Cerebello-Occipito-Parietal Module in the left hemisphere (Temporal Occipital Cortex (TempOcc), Extrastriate Visual Cortex (ExStr), and Cerebellum lobule VI), four nodes in the Prefrontal Module (two hubs in the right Dorsal Prefrontal Cortex (PFCd) and Lateral Prefrontal Cortex (PFCl) from both hemispheres), and one hub in sensorimotor module (left dorsal anterior cingulate cortex (dACC). The *Place* condition produced seven eigenvector hubs (Fig. 9B), located primarily in the Prefrontal Module (PFCd, PFCl, and medial posterior Prefrontal Cortex (PFRCmp) in both hemispheres, and one left Occipital hub in the Cerebello-Occipito-Parietal Module (TempOcc). Overall, the Grasp and *Place* network shared several hubs (TempOcc, PFCd, and PFCl), but the rest were distinct. Specifically, the grasp network had more occipital and cerebellar hubs (from Module 1), and the place network had more frontal hubs (from Module 2).

**Figure 9.**
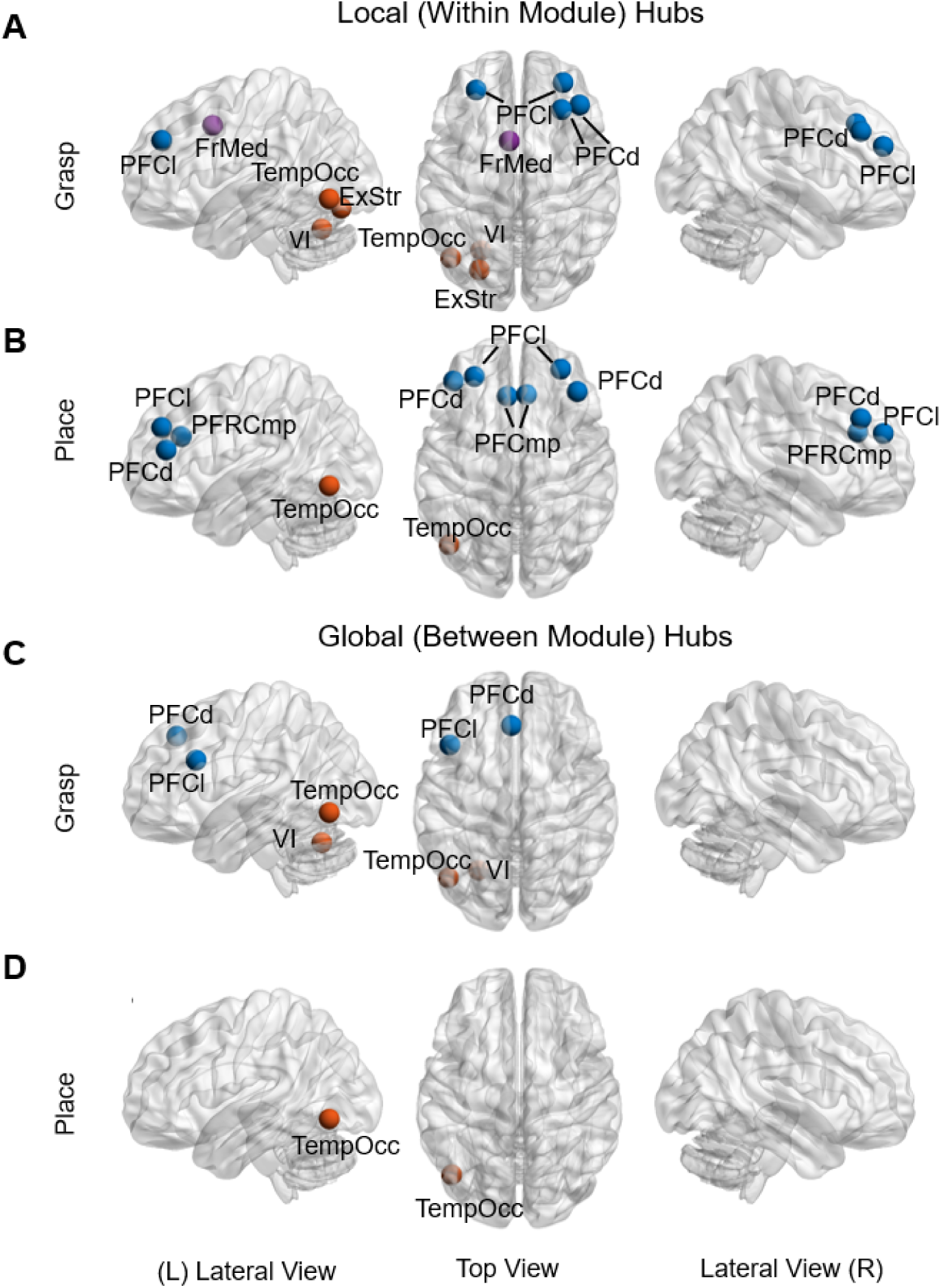
Brain maps showing local and global hubs for the *Grasp* and *Place* conditions. Hubs were defined as nodes at the 95^th^ percentile or higher and were significantly different from null models at alpha < 0.05 FDR-corrected. Local hubs are calculated using the eigenvector centrality for *Grasp* (A) and for *Place* (B). Global hubs are calculated using the betweenness centrality for *Grasp* (C) and for *Place* (D). Blue nodes are part of the Prefrontal Module, and red nodes are part of the Cerebello-Occipito-Parietal Module. Abbreviations: ExStr: Extrastriate Visual Cortex, ParOcc: Parietal Occipital Cortex, PFCd: Dorsal Prefrontal Cortex, PFCl: Lateral Prefrontal Cortex, PFClv: Ventrolateral Prefrontal Cortex, PFCmp: medial posterior Prefrontal Cortex, TempOcc: Temporal Occipital Cortex, TempPole: Temporal Pole, VI: left Cerebellum, VIII: Left lobule VIII Cerebellum, Crus I: Left Crus I Cerebellum

We used *betweenness* centrality to identify nodes that were highly correlated across modules. In the grasp network (Fig. 9C), we identified 2 global hubs in the Prefrontal Module (PFCd and PFCl in the left hemisphere) and 2 global hubs in the Cerebello-Occipito-Parietal Module (TempOcc and Cerebellum lobule VI in the left hemisphere). In contrast, the place network only had a global hub in left TempOcc, which is part of the Cerebello-Occipito-Parietal Module (Fig. 9D). Overall, *grasp* and *place* share a unique global hub in the temporal occipital cortex, but *grasp* had a greater number of nodes identified as hubs.

## DISCUSSION

Previous studies of movement control have largely focused on the initial act of prehension (i.e., Desmurget et al., 1996; Grafton, 2010; Karl & Whishaw, 2013), but less is known about the object placement that is often the ultimate goal of the interaction. In particular, it is not known how the intention to reach or place, and the resulting differences in sensory feedback, influence cerebello-cortical activation and functional networks. To address these questions, we conducted an alternating reach-to-grasp and reach-to-place experiment, compared the associated BOLD activation at the whole brain and ROI levels, used the latter to decode task type, and investigated functional connectivity metrics using GTA. Given that both actions involved the same objects and transport kinematics, we expected to see an overlap in the brain regions activated, but hypothesized that differences in the pattern of brain activity and cerebello-cortical functional connectivity would emerge due to the differences in intention and sensory feedback. Our results confirm that different structures were recruited (especially in the cerebellum), that action-related ROIs were modulated by task, and that functional network modularity was task-dependent.

### Univariate BOLD Analysis during Planning and Execution

To understand the relationship between grasp and placement control, we first performed a whole-brain analysis across movement preparation and execution. In the following sections, we consider first their commonalities, then their differences.

#### Common (Task-independent) Cortical and Subcortical Activation

*Planning Phase*. Our data show that *grasp* and *place* elicit a similar widespread increase in brain activation in preparation for action, primarily in the bilateral frontal, temporal, and occipital cortices, as well as in the left cerebellum, and to a specific parietal activation in the inferior parietal lobule. While extensive cortical activation during action planning has been reported in previous work (Baltaretu et al., 2020; Cappadocia et al., 2017), we also observed extensive cerebellar activation. Consistent with ipsilateral (to the right hand) lateralization (Grodd et al., 2001; Mottolese et al., 2013), we observed activation in the right Crus I and lobule VI, which have been implicated in bridging higher-order planning and motor control (Chabrol et al., 2019; Gao et al., 2018; Koziol et al., 2014; Park et al., 2018; Sokolov et al., 2017; Wagner et al., 2019; Zhu et al., 2023). We also saw a widespread left cerebellar activation linked to visuospatial processing (Starowicz-Filip et al., 2021), which was generally similar in both tasks.

*Execution Phase*. During movement, premotor cortex, SMA, SPL, SPOC, and LOC were recruited in both tasks, presumably to maintain movement goals and correct movement errors (Cappadocia et al., 2017; Filimon, 2010). Finally, there was extensive bilateral cerebellar involvement. This bilateral action could support communication between the left and right cerebellum, as visuospatial information is shared with the right hemisphere, which is then recruited for ipsilateral motor control. These findings represent a general reach transport-driven commands, feedback, and adjustments that do not dissociate the specific movement type.

Furthermore, there is a general shift in activation between these two phases as one progresses from planning the action to its execution, with planning showing greater involvement of PFC and IPL, and execution showing greater recruitment of sensorimotor regions and SPL. This reflects a transition from conscious motor intention preparation, as the movement goals are taken into account, to the integration of somatosensory cues and achieve the motor output (Castiello et al., 2000; Desmurget & Sirigu, 2012; Gamberini et al., 2021). This temporal distinction could hint at a processing differentiation in the parietal cortex as the action is planned and executed.

#### Selective (Task-dependent) Cortical and Subcortical Activation

*Planning Phase*. A whole brain voxelwise ‘*grasp* vs *place’* contrast revealed an increase in activation in the left (occipital) lingual gyrus for *grasp* and two cerebellum regions (right lobule V and VIIIa) for *placement*. As part of the ventral stream, the lingual gyrus is thought to be involved in identifying objects’ features required to perform the *grasp* (Kumral et al., 2025; Palejwala et al., 2021; Valyear & Culham, 2010). Respectively, cerebellar lobules V and VIIIa have been shown to implement visual working memory and the process of proprioceptive information for motor control (Bhanpuri et al., 2013; Boisgontier & Swinnen, 2014; Brissenden et al., 2021; Rondi-Reig et al., 2014).

These placement-specific activations suggest a role of the cerebellum in processing visual information to compare the object to the template target’s features for a placement-specific motor plan, also taking into account the body’s position when holding the object (Boisgontier & Swinnen, 2014; Therrien & Bastian, 2015). Presumably, this involves thalamic connections linking the ipsilateral cerebellum to contralateral cortex (Spampinato et al., 2024; Therrien & Bastian, 2015).

Finally, we observed greater preparatory activation in primary motor and somatosensory cortex during placement (Fig. 6A). Previous studies have shown that somatomotor cortex activity can predict planned action (Ariani et al., 2022; Gale et al., 2021). Our work shows that this activity also predicts the action type. Participants held a block at the start of both tasks (see METHODS), so these differences cannot be due to passive sensory feedback, but reflect the attentional gating of that feedback (since the initial block was only relevant in the *place* task) and/or different movement intentions.

*Execution Phase*. During action, we observed a relative increase in BOLD response in occipital regions (cuneal cortex, lingual cortex, and occipital fusiform gyrus) and the cerebellum during *placement*. Increased occipital activation could reflect the more complex visual feedback required to exactly match block location and orientation to the template (Luabeya et al., 2024). This can also be viewed as comparing a target to a landmark, which is known to engage ventral occipital cortex (Chen et al., 2014; Musa et al., 2025). In contrast, our *grasp* task was more forgiving, i.e., grip variations could be corrected after contact with the object (Lederman & Wing, 2003; Lukos et al., 2007; Paulun et al., 2014; Rearick & Santello, 2002).

#### Region of Interest Analysis and Classification

Although our voxelwise analysis suggested task overlap across large areas of cortex, this does not preclude the possibility of more subtle modulations within this activation. To test this possibility, we performed an ROI analysis of several well-established areas for visually-guided action (Castiello & Begliomini, 2008; Culham et al., 2003; Culham & Kanwisher, 2001; Monaco et al., 2024). We did not find any differences during *execution*, but we found several significant task differences during the *planning* phase, with placement showing greater activation in aIPS, M1, S1, and PMd. Again, these differences might be accounted for by both the need to process somatosensory information from the block and more complex visuospatial relations in the *Place* task. Finally, our classification analysis confirmed that activity in these regions differentiated between our *Grasp* vs*. Place* tasks, a result that may be relevant for the application of brain-machine interfaces for real-world reach-place sequences (Lebedev & Nicolelis, 2017; Nicolas-Alonso & Gomez-Gil, 2012).

Canonically, aIPS has been described as important for computing object features relevant to grasping (Frey et al., 2005; Rice et al., 2006). Indeed, compared with less fine-tuned actions, such as reaching or pointing, grasp exhibits greater activation in this parietal region. However, when it is contrasted with another action requiring a comparable level of motor precision, we found greater recruitment of aIPS for placement. Demonstrating that aIPS is not an exclusively grasp-related region but carries object features relevant for dexterous motor control. Its greater activation during placement could reflect an increased need for fine-tuning, given its reduced tolerance for movement errors. Perhaps placing an object in a wider space, which does not require a highly precise movement, would not elicit such an increase in aIPS involvement. Similar to the difference observed between performing a precision grasp on a small object and a power grasp on a larger object, which does not need the same intricacy in the movement (Cavina-Pratesi et al., 2018; Ehrsson et al., 2000). Therefore, aIPS involvement goes beyond a finger configuration for grasping; instead, it gathers object features relevant to precise maneuvering for successful object interactions.

### Graph Theoretical Functional Connectivity Analysis

To understand the similarities and differences between grasp and placement at a functional network level, we used a graph theoretical approach. This allowed us to assess the distribution of correlated BOLD time series at three levels: global (macro-level), intermediate (meso-level), and local (micro-level). This analysis again revealed both functional similarities and differences between our two motor tasks, as discussed below.

#### Macro-scale Network Features: Equivalence of Global Parameters

At the macro-scale, global GTA parameters suggest that grasping and placing mechanisms share similar levels of information transfer efficiency, overall segregation, and synchrony. This may reflect the fact that in both cases the brain was in a similar ‘motor control’ state (Madole et al., 2023), related to the shared transport mechanisms discussed above. However, it could also be the case that global measures are simply not sufficiently specific to distinguish between very similar behaviors (Di et al., 2013; Ghaderi et al., 2023b; Luabeya et al., 2026; Musa et al., 2025).

#### Meso-scale Network Features: Similar Modules, Different Community Strengths

Our modularity analysis revealed the same subnetworks in both tasks: a cerebello-occipito-parietal module, a prefrontal module, and a sensorimotor module. This segregation likely reflects three major processes: 1) higher-order cognitive processes (such as the assimilation of the instruction into the motor plan in the prefrontal module, 2) incorporation of visuospatial cues into the motor plan through the cerebello-occipito-parietal module, and 3) action control in the sensorimotor module (Luabeya et al., 2026). We did not observe segregation of dorsal-ventral visual streams previously documented in a simple pointing task (Musa et al., 2025), possibly because the current tasks require greater processing of both object location and orientation information (Monaco et al., 2011, 2020; Velji Ibrahim et al., 2022). Finally, the modularity scores of the sensorimotor module were similar in both tasks, likely because they both place high demands on somatomotor cortex.

However, the modularity scores of the other two modules showed significant task differences (Fig. 7). G*rasp* showed higher modularity in the cerebello-occipital-parietal module, whereas *place* showed higher modularity in the prefrontal module. This suggests that the prefrontal module had less functional connectivity with other modules in the place task. This shift in neural topology demonstrates selective task-specific intra-module strengthening (Finc et al., 2026) and is consistent with the lack of prefrontal ‘betweenness hubs’ in this task (see the next section). Conversely, this suggests that the cerebello-occipital-parietal module functioned more independently in the grasp task.

Finally, omitting the overall trend in the data (the general rising trend of BOLD activation across the entire brain during movement planning and execution) affected the *place* condition more than the *grasp* condition (Fig. 8). Specifically, this resulted in a division of the cerebello-occipito-parietal module into two submodules: a highly modular occipito-parietal module and a less modular cerebellar module. Previous work has suggested that global trends in the data sustain distal communication, such that trend removal highlights more local processes (Musa et al., 2025). This is consistent with the notion that the cerebello-occipito-parieto module is involved in the integration of multimodal information in both our tasks.

#### Micro-scale Network Features: Local vs. Global Hubs

Because GTA measures are based on correlations between nodes, important hubs may not necessarily correspond to the nodes with the highest activation (Tomasi et al., 2014). However, our hubs (Fig. 9) fell within regions that showed increased activation (Figs. 3 and 4), suggesting that highly activated areas tend to also share signals broadly with other regions. Once again, at this scale, we found both commonalities and differences between tasks.

Both tasks shared a temporal occipital hub associated with the dorsal attention network (Schaefer et al., 2018) that served as both a local (within-module) and a global (between-module) hub. Multi-centrality hubs, which are important for both local and global connectivity, are particularly robust and essential in the network topology (Bertolero et al., 2017; Oldham et al., 2019). Temporal occipital cortex, part of the ventral visual stream (Goodale & Milner, 1992), is involved in both object recognition and object-object spatial relations (Avanzini et al., 2013; Chen et al., 2014), which play a role in both tasks (i.e., hand-object / object-template recognition and matching). It is thus expected that this region would show both internal and global connectivity for both tasks.

In addition, *grasp* and *place* shared local hubs in the lateral and dorsal prefrontal cortex, parts of the ventral attention network, and the fronto-parietal control network (Schaefer et al., 2018). These regions are associated with motor control inhibition, information value processing, and response selection during tasks (Dixon & Christoff, 2014; Krämer et al., 2013; Rowe et al., 2000). Overall, these hubs reflect the general cognitive, sensory, and motor demands of both tasks

Finally, we also found task-specific differences in both local and global hubs. In terms of significant local hubs, the *place* task had more eigenvector hubs in the prefrontal cortex than the *grasp* task (6 vs. 4), re-emphasizing the internal computations in areas such as PFCmp and PFCd, which are involved in motor correction after error (Danielmeier et al., 2011). This might reflect a greater need for accuracy in the place task. In contrast, the grasp task showed an additional hub in the sensorimotor module, specifically, in dorsal anterior cingulate cortex (’FrMed’), an area associated with error prediction (Modirrousta & Fellows, 2008; Schulz et al., 2011). Grasp also showed more hubs in the cerebello-occipito-parietal module. Together, these might reflect the need for more local computations for feed-forward control in the *grasp* condition.

In terms of global connectivity, *grasp* showed betweenness hubs in prefrontal cortex (PFCd and PFCl), the temporal occipital region, and cerebellar lobule VI, being involved in the long-range communication. These might reflect stable connections underlying well-established internal models of the world required for feed-forward control (McNamee & Wolpert, 2019; Wolpert et al., 1995). In contrast, *placement* showed fewer betweenness hubs overall, indicating a less integrated system, perhaps reflecting less reliance on internal models and more reliance on sensory feedback. In particular, placement showed no significant prefrontal hubs, consistent with the increase in modularity discussed above. However, it maintained the temporal occipital hub, perhaps reflecting communication of visual object-template relationship with the sensorimotor module (Musa et al., 2025).

### Limitations and Future Direction

We acknowledge several limitations that may influence the interpretation of our results. First, even though the runs in the study were pseudorandomized, there could be an order-related variable, as the experimental design required alternating between grasp and place. Whereas this was partially mitigated by our counterbalanced order of runs, within a run, a grasp was still followed by a placement, and a placement by a grasp. Therefore, a residual activity might be carried over from one trial to the next and obscure the differentiation between our conditions (Aguirre, 2007; Buckner, 1998). While a long intertrial interval was used to give sufficient time for the hemodynamic response to return to baseline, it is still possible that some residual noise was carried from the previous trial. Therefore, the differentiation between our tasks may not have been fully captured; however, we were still successful in demonstrating a noticeable difference between grasp and placement. Since placement required the presence of an object held in hand and grasp does not, even without order-dependency, participants would still be aware of the action they will perform in the upcoming trial. To address some aspects of the task predictability, we varied the extrinsic object features (orientation and location). Future work could also include incongruent tasks in which the action is planned but not executed.

Second, looking more specifically at the GTA analysis, because of the nature of an event-related task, our analysis was executed on a relatively short window from which the correlation was extracted. This could result in unstable trial-by-trial edge estimates. Therefore, we avoided making assumptions based on individual trials, aggregated their outputs within subjects, and conducted network-wide analysis to understand the system as a whole. Furthermore, we compared our GTA measures to a null model and conducted cross-validation to substantiate the robustness of our findings. Other neuroimaging studies have successfully applied GTA and functional connectivity analyses on shorter event-related studies (Luabeya et al., 2026; Musa et al., 2025; Shirer et al., 2012; Tomou et al., 2025). For future directions, one could address temporal constraints in a more naturalistic setting by repeating these experiments with electroencephalography (EEG), magnetoencephalography (MEG), or Near-Infrared Spectroscopy (fNIRS).

Finally, another limitation of the GTA modularity analysis stems from the implementation of its standard resolution parameter (Bassett et al., 2013), which limits the ability to identify more stringent communities in the network. Indeed, small modules may be less affected by detrending, as they might already account for the drift in the trial. Therefore, it would be less influenced by trend removal and better represent local communications. Future applications of GTA to sensorimotor control could optimize the resolution parameters used and improve module detection sensitivity.

### General Implications for Sensorimotor Imaging Studies

Manual control supports most of our interactions with the external world. Although the field of sensorimotor neuroscience has grown immeasurably through the study of simple reach-to-grasp movements, it is equally important to go ‘beyond reach’ to study how objects are used once they are grasped (Binkofski et al., 1999; Talati et al., 2005; Turesky et al., 2018). This particular study demonstrates that reach-to-grasp and reach-to-place movements, while superficially similar, evoke different neural mechanisms, likely due to the different computational requirements of these tasks (Luabeya et al., 2024; Lukos et al., 2007; Naish et al., 2013; Paulun et al., 2014; Rearick & Santello, 2002). A second outcome of this experiment was that neuroimaging studies tend to emphasize cerebral cortex, but in this study, the cerebellum proved important for distinguishing between our tasks. It is well known that this and other subcortical regions are crucial components in movement control (De La Peña et al., 2020; Ji et al., 2019), and their damage can result in drastic impairments (ie, Schucht et al., 2013; Wong et al., 2013), so these areas should be considered more in neuroimaging studies. Finally, we hope that this and similar studies (Luabeya et al., 2026; Musa et al., 2025) demonstrate the utility and importance of applying tools like GTA (often employed for the study of resting-state data in block designs) to event-related, behavioral paradigms. This approach provides a direct way to quantify global signal distributions, network modularity, and local network hubs in the context of the behavior that they have evolved to produce.

## Conclusion

This study addresses fundamental differences in whole brain activation and functional connectivity between reach-to-grasp and reach-to-place movements. Our results suggest that these behaviors share substantial commonality in their patterns of brain activation. However, they show meaningful differentiation in the importance of cerebellum in processing and maintaining the sensory information during the planning and execution of placement movements, in the decoding capacity of signals from action-related regions (aIPs, PMd, M1, and S1) during movement planning, and in the differentiating modularity between grasp and place. These differences could have real-world applications for understanding how brain damage affects the different components of manual interaction, and for brain-machine applications in realistic behavioral situations.

## SUPPLEMENTARY FIGURE

**Supplementary Figure 1.**
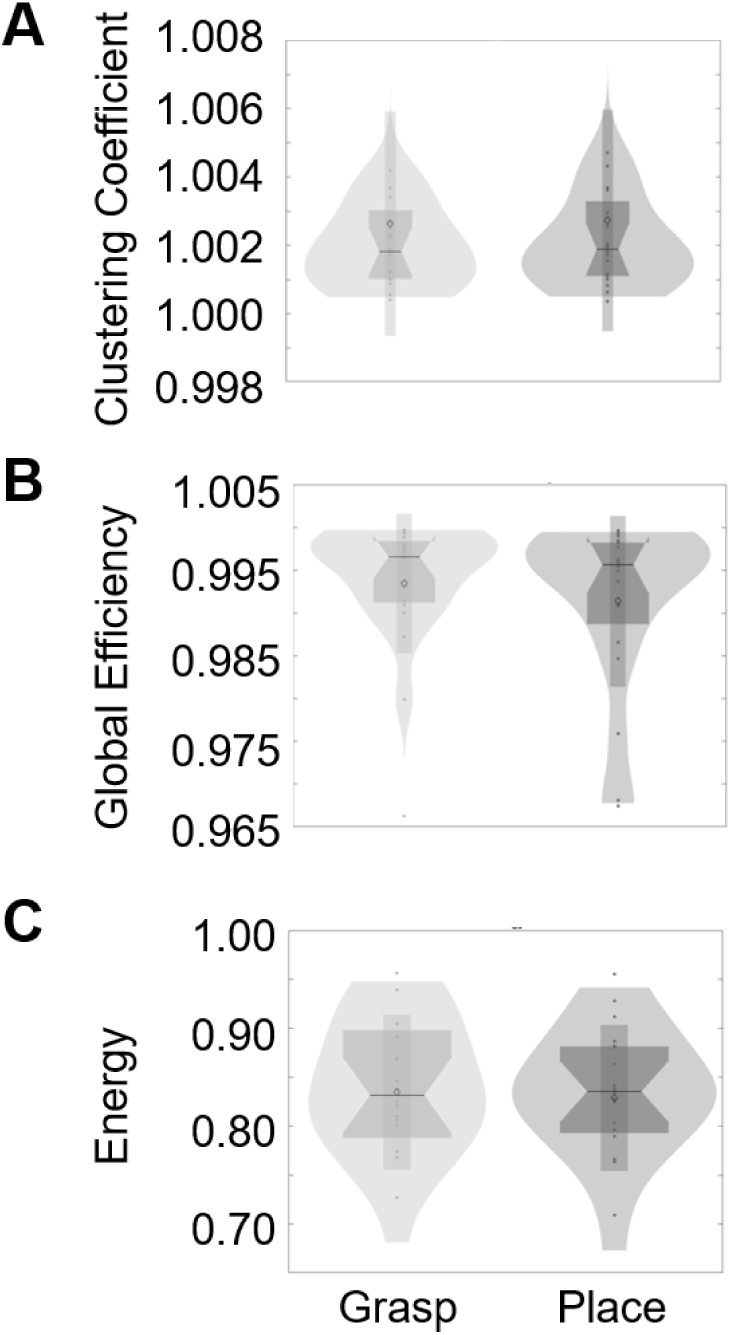
Violin plots displaying distributions of normalized global network parameters: clustering coefficient (A), global efficiency (B), and energy (C). To normalize these global measures, they were divided by the global measures outputs obtained from a null model. In all three cases, our nonparametric test did not find any significant difference based on task (*grasp* vs *place*).

## Article Information

### Data and Code Availability statement

Data and code are available for requests that comply with federal data sharing policies, pending approval from the requesting researcher’s local ethics committee and agreement on how to share author credits on any resulting publication.

### CRediT statement

GNL: Methodology, Investigation, Data curation, Formal Analysis, Conceptualization, Visualization, Writing - Original draft, Writing - review and editing. EF: Conceptualization, Methodology, Writing - review and editing. JDC: Conceptualization, Methodology, Writing - review and editing, Funding acquisition, Project administration, Supervision.

### Funding statement

This work was supported by grants from the National Science and Engineering Research Council (NSERC Grant RGPIN-2022-04527). Luabeya was supported by the Vision: Science to Applications Program, funded in part by the Canada First Research Excellence (CFREF) program. Crawford was supported by a Canada Research Chairs.

### Conflict of interest disclosure

the authors have none to report

### Ethics approval statement

This experiment was approved by the York University Human Participants Review Committee (Certificate #: e2023-336; initially approved November 09, 2023, and renewed November 02, 2025).

## Acknowledgments

The authors thank Dr. Xiaogang Yan, Dr. Diana Gorbet, Saihong Sun, and Christopher Giverin for technical support and Dr. Vishal Bharmauria, Dr. Jessica Parker, and Petros Geordiadis for comments on the manuscript.

## Notes

### Competing Interest Statement

The authors have declared no competing interest.

